# Alzheimer’s disease induced neurons bearing *PSEN1* mutations exhibit reduced excitability

**DOI:** 10.1101/2024.03.22.586207

**Authors:** Simon Maksour, Rocio K. Finol-Urdaneta, Amy J. Hulme, Mauricio Castro Cabral-da-Silva, Helena Targa Dias Anastacio, Rachelle Balez, Tracey Berg, Calista Turner, Sonia Sanz Muñoz, Martin Engel, Predrag Kalajdzic, Leszek Lisowski, Kuldip Sidhu, Perminder S. Sachdev, Mirella Dottori, Lezanne Ooi

**Affiliations:** School of Chemistry and Molecular Bioscience and Molecular Horizons, University of Wollongong, NSW, Australia; School of Medical and Indigenous Health Science and Molecular Horizons, University of Wollongong, NSW, Australia; Translational Vectorology Research Unit, Children’s Medical Research Institute, Faculty of Medicine and Health, The University of Sydney, Westmead, Australia; Australian Genome Therapeutics Centre, Children’s Medical Research Institute and Sydney Children’s Hospitals Network, Westmead, NSW 2145, Australia; Laboratory of Molecular Oncology and Innovative Therapies, Military Institute of Medicine - National Research Institute, Warsaw, Poland; Centre for Healthy Brain Ageing, School of Clinical Medicine, University of New South Wales, Sydney, Australia

**Keywords:** Alzheimer’s disease, *PSEN1*, neuronal excitability, iNs, iPSCs

## Abstract

Alzheimer’s disease (AD) is a devastating neurodegenerative condition that affects memory and cognition, characterized by neuronal loss and currently lacking a cure. Mutations in *PSEN1* (Presenilin 1) are among the most common causes of early-onset familial AD (fAD). While changes in neuronal excitability are believed to be early indicators of AD progression, the link between *PSEN1* mutations and neuronal excitability remains to be fully elucidated. This study examined induced pluripotent stem cell (iPSC)-derived NGN2 induced neurons (iNs) from fAD patients with *PSEN1* mutations S290C or A246E, alongside CRISPR-corrected isogenic cell lines, to investigate early changes in excitability. Electrophysiological profiling revealed reduced excitability in both *PSEN1* mutant iNs compared to their isogenic controls. Neurons bearing S290C and A246E mutations exhibited divergent passive membrane properties compared to isogenic controls, suggesting distinct effects of *PSEN1* mutations on neuronal excitability. Additionally, both *PSEN1* backgrounds exhibited higher current density of voltage-gated potassium (Kv) channels relative to their isogenic iNs, while displaying comparable voltage-gated sodium (Nav) channel current density. This suggests that the Nav/Kv imbalance contributes to impaired neuronal firing in fAD iNs. Deciphering these early cellular and molecular changes in AD is crucial for understanding the disease pathogenesis.

## 1 Introduction

Alzheimer’s disease (AD) is a devastating, progressive neurodegenerative disease that affects memory and cognition and is characterized by the loss of neurons. Research has shifted to looking at how early molecular changes may influence the cause and progression of AD. These studies focus on early changes in gene expression (Guennewig *et al*. 2021), energy metabolism (Johnson *et al*. 2020), altered neurogenesis and neuronal differentiation (Meyer *et al*. 2019; Arber *et al*. 2021), and neuronal firing (Spoleti *et al*. 2022; Ghatak *et al*. 2019). Changes in neuronal excitability have been predicted to be an early predictive and driving force of neurodegeneration and disease pathology, in different diseases, including AD (as reviewed in Targa Dias Anastacio *et al*. (2022)) and amyotrophic lateral sclerosis (as reviewed in Do-Ha *et al*. (2018)). Neuronal excitability changes in disease are influenced by both Aβ (Busche *et al*. 2008; Busche *et al*. 2012) and tau pathology (Busche *et al*. 2019; Crimins *et al*. 2012), whilst conversely, excitability changes can also drive both AD hallmark pathologies (Cirrito *et al*. 2005; Yamamoto *et al*. 2015; Pooler *et al*. 2013; Wu *et al*. 2016). Futhermore, there are demonstrated contributions from inhibitory neurons (Verret *et al*. 2012; Ghatak *et al*. 2019; Nuriel *et al*. 2017) and glial cells (Targa Dias Anastacio *et al*. 2022). Correcting neuronal activity through pharmacological or genetic intervention in AD mouse models improves memory and cognition (Martinez-Losa *et al*. 2018; Roberson *et al*. 2007; Ping *et al*. 2015), highlighting the role of neuronal excitability regulation in disease progression.

*PSEN1* is the most common causative gene for early-onset AD (also known as familial AD, fAD) and is believed to contribute to neuronal vulnerability through the overproduction of toxic amyloid-ß (Aβ) peptides, which results in the generation of Aß plaques (Ooi *et al*. 2020). There are over 300 known mutations in *PSEN1*, many with pathogenic outcomes, however the effects of each mutation on the disease phenotype remains to be fully elucidated (*PSEN1* mutations database, ALZforum. https://www.alzforum.org/mutations/psen-1). In addition to disrupted amyloid precursor protein (APP) processing and plaque formation, *PSEN1* mutations induce early changes in neurons, including, increased susceptibility to Aβ (Armijo *et al*. 2017) and ferroptosis (Greenough *et al*. 2022), dysregulated neurogenesis and differentiation (Arber *et al*. 2021; Hurley *et al*. 2023), decreased neurite outgrowth (Dowjat *et al*. 1999; Furukawa *et al*. 1998; Balez *et al*. 2016; Ghatak *et al*. 2019), endosomal dysfunction (Kwart *et al*. 2019) and neuronal excitability (Hurley *et al*. 2023; Vitale *et al*. 2021; Chen *et al*. 2021). These studies highlight the complex role of presenilin-1 (PSEN1) plays in multiple cellular, and the need to understand how specific mutations affect neuronal function.

Human pluripotent stem cells (iPSCs) offer an avenue to generate neurons from patients bearing a disease-relevant gene mutations, and interrogate the intrinsic excitability properties of neurons in the absence of late-stage AD pathology and supporting cell types. Therefore, this study aimed to use iPSC-derived NGN2 induced neurons (iNs) from two AD patients bearing either a S290C or A246E mutation in *PSEN1*, along with their CRISPR-corrected isogenic controls to investigate early neuronal excitability changes in disease. Understanding the common and divergent cellular and molecular changes disrupted in neurons from AD patients bearing different *PSEN1* mutations will provide insight into the cause and progression of neurodegeneration, and allow for targeted therapeutic intervention.

## 2 Methods

### 2.1 Cell culture

#### 2.1.1 iPSC cell lines and maintenance

Use of iPSC lines for this project was approved by the UOW Human Ethics Committee (#2017-375, 2017-382, 2020-450, 2020-451, 13-299). This study used iPSCs generated from early-onset AD patients with a *PSEN1* S290C (S290C) (**Figure S1**) or a A246E mutation (A246E)(Muñoz *et al*. 2018) and their respective CRISPR-corrected isogenic controls, S290^IC^ (**Figure S1**) and A246^IC^ (Targa Dias Anastacio *et al*., 2024, in revision). The iPSCs were cultured as previously reported in Abu-Bonsrah *et al*. (2019), Denham and Dottori (2011) and Mattei *et al*. (2019). Briefly, the iPSCs were maintained in mTesR1 (StemCell Technologies, #85850) on matrigel-coated tissue culture ware, kept in normoxic conditions at 37 °C with 5% CO_2_. Cells were passaged once every 5-7 days using 0.5 mM EDTA (Life Technologies, #AM9260G) in PBS^-/-^ (Life Technologies, #14190250). Methods on the cell line generation and characterization can be found in the Supplementary materials.

### 2.2 Lentiviral production

Viral particles containing an open reading frame of *Neurogenin-2* (*NGN2*) were produced to differentiate iPSCs into mature neurons as described in Hulme *et al*. (2020) and Maksour *et al*. (2024). Briefly, HEK293T cells were transfected with the DNA of lentiviral packaging plasmids vSVG (Addgene, USA, #8454), RSV (Addgene, #12253), pMDL (Addgene, #12251), and either the tetracycline transactivator (TTA) vector, M2rtTA (Addgene, #20342), or the *NGN2* overexpression vector, TetO-*NGN2*-eGFP-Puro plasmid (Addgene, #79823) using Polyethyleneimine (Sigma-Aldrich, USA, #408727). DNA was added in a ratio of 4:2:1:1, transfer vector:pMDL:RSV:vSVG. The cell culture media containing viral particles was collected every 24 hours over 3 days. The viral supernatant was concentrated 200x by ultracentrifugation at 66,000 x *g* for 2 h at 4 °C. The viral pellet was resuspended in PBS and stored at −80 °C until needed.

### 2.3 Generation of NGN2*-*induced neurons (iNs)

This study used a protocol published in Maksour *et al*. (2024) to generate mature neurons via NGN2 overexpression. Briefly, iPSCs were resuspended as single cells using Accutase for 2-3 min at RT. Single cells were plated at 15,000 cells/cm^2^ onto 10 μg/mL poly-D-lysine (PDL) and laminin (LAM) coated culture plate in mTeSR1 media supplemented with 10 μM Y27632. Cells were allowed to attach for 6-8 hours, after which 0.5 μL of viral particles of both NGN2 overexpression and the TTA per 15,000 cells. Virus was removed 16-20 hours following transduction with fresh neural media [Neurobasal medium (NBM; Life Technologies, #21103-049) supplemented with 1x N-2 supplement 1x B-27 supplement, 1x Insulin-Transferrin-Selenium-A and 2 mM L-glutamine] supplemented with 1 μg/mL doxycycline (DOX; Sigma-Aldrich, # D9891), 10 μM SB431542 and 0.1 μM LDN193189 to promote a cortical fate. After 24 hours of DOX induction, 0.5 μg/mL puromycin was added daily for 3 days for selection of successfully transduced cells, in addition to DOX, SB431542 and LDN193189. Following selection fresh media supplemented with 10 μg/mL BDNF was added. Following selection, BrainPhys media [Brainphys medium (StemCell Technologies, #05790) supplemented with NeuroCult SM1 (without vitamin A; StemCell Technologies, #05731) and N2 supplement-A (StemCell Technologies, #07152)] was subsequently added in at increasing concentrations (25-100% in NM) for each media change to improve maturation. Neurons were assessed by whole-cell patch clamp between 21-35 days post viral transduction.

### 2.4 Electrophysiology

Sterile plastic coverslips, cut into 10 mm slides, were coated with PDL and LAM before plating NGN2 iNs for functional characterization. Whole-cell patch clamp recordings were performed on matured neurons aged 3-5 weeks, following the protocol outlined in Hulme *et al*. (2020). Recordings were conducted at room temperature (20-22ºC) using a MultiClamp 700B Amplifier, digitized with a Digidata 1440, and controlled via pClamp11 software (Molecular Devices). Whole-cell membrane currents were measured at 100 kHz, with series resistance compensated at 60-80%. Fire-polished borosilicate patch pipettes with a resistance of 2-4 MΩ were employed with an intracellular buffer composed of (in mM) 140 K-gluconate, 10 NaCl, 2 MgCl_2_, 10 HEPES, 5 EGTA (pH 7.2, osmolality 295 ± 5 mOsm/kg). The bath solution for current clamp experiments contained (in mM) 135 NaCl, 2 CaCl_2_, 2 MgCl_2_, 5 KCl, 10 glucose, 10 HEPES (pH 7.4, osmolality 315 ± 5 mOsm/kg). The resting membrane potential (RMP) was determined as the average membrane voltage without current injection, while the rheobase was defined as the minimum current needed to elicit a single action potential.

## 3 Results

### 3.1 iPSC derived neurons from two different fAD *PSEN1* backgrounds exhibit impaired excitability

We have previously demonstrated the forced overexpression of NGN2 successfully and robustly generates functional, glutamatergic excitatory induced neurons (iNs) (Maksour *et al*. 2024). This study generated iNs from iPSCs from fAD patients bearing the *PSEN1*^S290C^ (S290C) and *PSEN1*^A246E^ (A246E) and their respective CRISPR corrected counterparts, S290^IC^ and A246^IC^, to assess their electrophysiological properties (**Figure 1A**). Neuronal excitability was evaluated by manual patch-clamp in the whole-cell configuration with representative recordings displayed in **Figure 1B** with data for all parameters summarized in **Table S2**.

**Figure 1.**
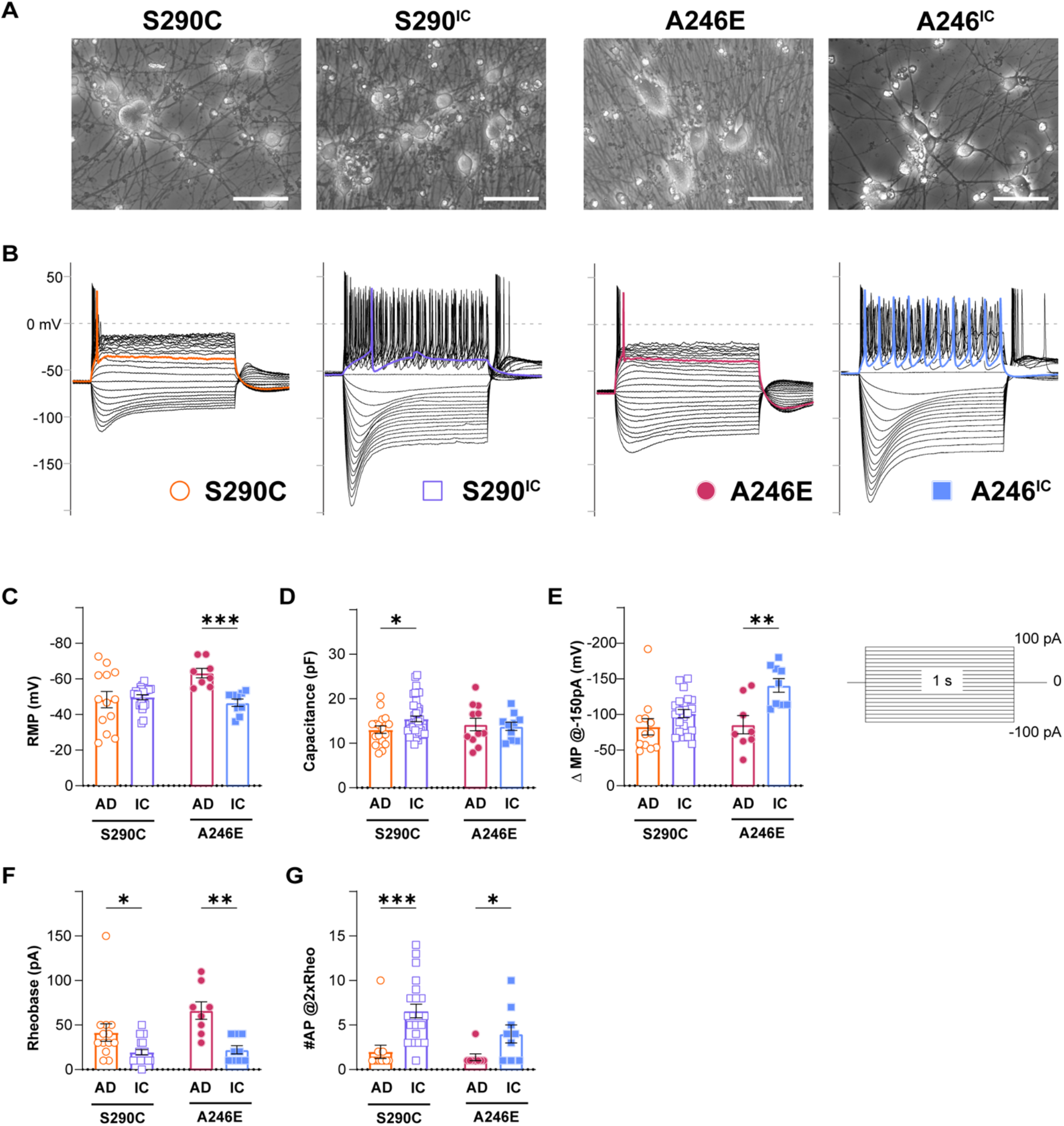
Neurons derived from AD patients bearing a *PSEN1* mutation exhibit reduced excitability. Whole cell patch clamping was performed on day 21-35 NGN2 iNs derived from fAD patients with a *PSEN1*^S290C^ (S290C) and *PSEN1*^A246E^ (A246E) and their respective CRISPR-corrected isogenic controls, S290^IC^ and A246^IC^ (**A**). **(B)** Representative current clamp recordings of membrane potential responses of all cell lines. The stimulation protocols consisted of one-second-long current injections from −100pA to 100pA in 10pA steps. The colored traces indicate firing activity at rheobase. Neurons were tested for resting membrane potential **(C)**, capacitance **(D)**, change in membrane potential **(E)**, rheobase **(F)** and the number of action potentials fired at 2 times rheobase **(F)**. Data is presented as the mean ± SEM. Each data point represents an individual cell (n = 7 – 41), from 3 independent differentiations. Data was analysed using multiple t-tests with Holm-Sidak for multiple comparisons where \**p* < 0.05, \*\**p* < 0.01, \*\*\**p* < 0.001 and \*\*\*\**p* < 0.0001. **Abbreviations**: AD – Alzheimer’s disease, IC – isogenic control.

The resting membrane potential (RMP) was evaluated within 2 minutes of switching to the current-clamp mode. RMP showed no difference between S290C and S290^IC^ iNs, while A246E iNs had significantly lower RMP compared to their isogenic controls (−63.3± 2.7 mV, n = 8 vs −46.6 ± 2.9 mV, n = 9, respectively; *p* < 0.001) (**Figure 1C**). S290C iNs had significantly lower capacitance than S290^IC^ (13.0 ± 0.8 mV, n = 17 vs 15.5 ± 0.6 mV, n = 41, respectively, *p* < 0.05) reflecting the generation of smaller iNs from this fAD line (**Figure 1D**), while A246E and A246^IC^ capacitance values were comparable.

iNs were stimulated through to 1 s long step current injections (−100 pA to −50 pA, **Figure 1**) under current-clamp. Negative current injections elicited hyperpolarizing membrane responses (sags) consistent with the activation of hyperpolarization-gated cyclic nucleotide-activated ion channels (HCN). Quantification of the change in membrane potential (ΔMP) evidenced upon −150 pA current injections revealed similar hyperpolarizing sags in S290C and S290^IC^ iNs, and larger sags in A246E iNs compared to its isogenic control (ΔMP = −85. 8 ± 12.8 mV, n = 8 vs −141.0 ± 9.5 mV, n = 9, *p* < 0.01) (**Figure 1E**).

The rheobase (the minimum current injection required to induce one action potential, AP) was higher in both fAD iNs (S290C: 41.5 ± 9.8 pA, n = 13; A246E: 66.3 ± 9.8 pA, n = 8) compared to their corrected controls S290^IC^ (19.5 ± 3.0 pA, n = 21; *p* < 0.05) and A246^IC^ (22.2 ± 4.7 pA, n = 10; *p* < 0.01), respectively (**Figure 1F**). Notably, positive current injections (up to 100 pA) elicited AP firing in all iNs, but multiple action potentials were only observed in corrected iNs (**Figure 1B**). Accordingly, the number of APs fired upon current injections equivalent to two times the rheobase (#AP@2xRheo) was significantly lower in both fAD lines compared to corrected iNs (**Figure 1G**). Thus, iNs derived from *PSEN1* S290C iPSCs fired 2.0 ± 0.7 action potentials (n = 12) compared to 6.6 ± 0.8 (n = 21) APs by S290^IC^ (*p* < 0.001), and *PSEN1* A246E fired 1.4 ± 0.4 APs (n = 8) compared to 4.0 ± 1.0 APs (n = 9) by A246^IC^ iNs (*p* < 0.05). Thus, neurons derived from fAD patients with S290C or A246E mutations in *PSEN1* exhibited reduced firing capacity and excitability compared to their isogenic controls, indicating a common phenotype of reduced neuronal excitability.

### 3.2 iNs from different fAD *PSEN1* backgrounds display distinct AP characteristics

Comparative analysis of AP shapes (at rheobase) between fAD and corrected iNs unveiled disparities in AP peak amplitude, area, half-width, rise time, rise slope, and maximal decay slope, as outlined in **Table S1**. While AP peak amplitude showed no variance between S290C and S290^IC^ neurons (**Figure 2A**), their AP area differed significantly, with S290C iNs exhibiting over a 5-fold smaller area than their isogenic control (2229.0 ± 423.5 mV*ms, n =13; vs 12036.2 ± 1390.3 mV*ms, n = 21; *p* < 0.001) (**Figure 2B**). Additionally, the AP rise slope was markedly higher in S290^IC^-derived neurons (59.3 ± 9.5 mV/ms, n = 21) compared to those harboring the S290C mutation (25.6 ± 4.1 mV/ms, n = 13, *p* < 0.05) (**Figure 2D**). Consequently, the AP rise time was more than 2-fold slower in S290C neurons compared to S290C^IC^ (1.3 ± 0.2 ms, n = 13 vs 0.6 ± 0.2 ms, n = 20, *p* < 0.05) (**Figure 2E**) with no discernible differences in AP maximal decay slope (*p* = 0.15) (**Figure 2F**).

**Figure 2.**
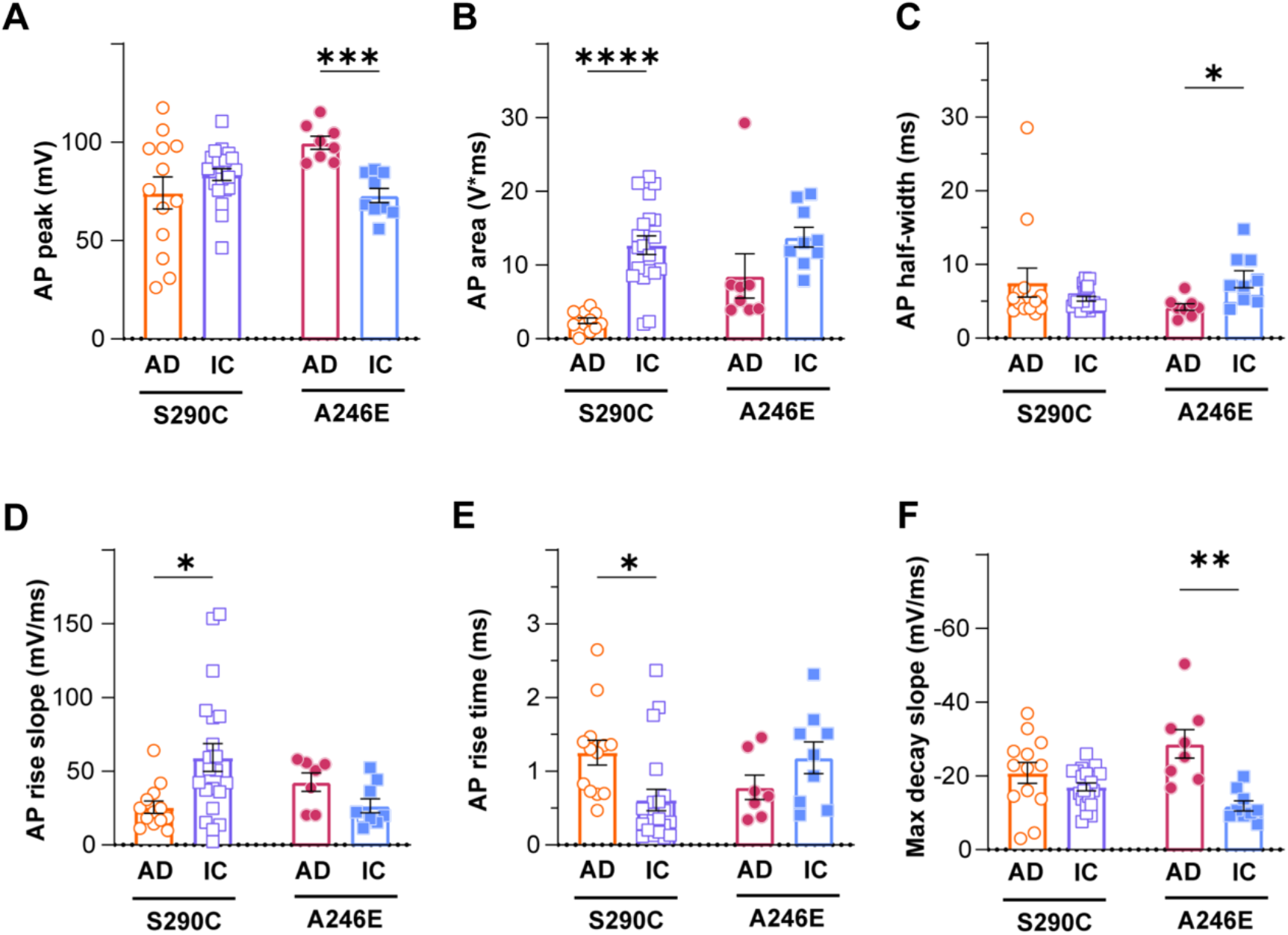
AD induced neurons display altered action potential properties compared to isogenic controls. Action potential properties were also compared between AD patient lines with *PSEN1* mutations (S290C and A246E) and isogenic controls, including peak amplitude **(A)**, area **(B)**, half-width **(C)**, rise slope **(D)**, rise time **(E)** and max decay slope **(F)**. Data is presented as the mean ± SEM. Each data point represents an individual cell (n = 7 – 41), from 3 independent differentiations. Data was analyzed using multiple t-tests with Holm-Sidak for multiple comparisons where \**p* < 0.05, \*\**p* < 0.01, \*\*\**p* < 0.001 and \*\*\*\**p* < 0.0001. **Abbreviations**: AD – Alzheimer’s disease, IC – isogenic control.

In contrast, PSEN1 A246E fAD iNs exhibited higher AP peak amplitudes (99.7 ± 3.3 mV, n = 8) compared to corrected A246^IC^ iNs (72.9 ± 3.7 mV, n = 9; *p* < 0.001) (**Figure 2A**), while showing no significant difference in AP area (*p* = 0.44) (**Figure 2B**). Despite similar AP rise parameters (**Figures 1D-E**), A246E neurons displayed narrower APs than their isogenic control (AP half-width 4.2 ± 0.5 ms, n = 8 vs 8.0 ± 1.2 ms, n = 9, respectively; *p* < 0.05) (**Figure 2C**), accompanied by markedly larger maximal decay slopes (−28.7 ± 3.8 mV/ms, n = 8 vs −11.9 ± 1.4 mV/ms, n = 9, *p* < 0.01)(**Figure 2F**). Thus, the action potential shape in fAD iNs appears distinctively affected by specific *PSEN1* mutations.

### 3.3 Voltage-gated potassium currents are larger in iNs from both *PSEN1* fAD patients

The shape of the action potential is intricately governed by factors such as ion channel dynamics and the equilibrium between inward and outward currents, primarily mediated by voltage-gated sodium (Nav) and potassium (Kv) channels, respectively. In **Figure 3A**, representative iN whole-cell currents elicited by 250 ms step depolarization (from −80 mV to 50 mV; Vh −80 mV, inset) recorded in voltage-clamp mode are displayed for fAD *PSEN1* mutations and their isogenic controls. Total Nav and Kv currents (at −10 mV and 20 mV, respectively) were normalized to the cell capacitance and expressed as current density (pA/pF). While no detectable differences in Nav channel-mediated currents were observed between disease lines and isogenic controls (**Figure 3B**), currents mediated by Kv channels were significantly larger in neurons derived from patients with fAD *PSEN1* mutations (S290C: 161.5 ± 28.1 pA/pF, n = 17; A246E: 89.2 ± 7.1 pA/pF, n = 10) compared to their respective CRISPR-corrected isogenic controls (S290^IC^: 97.0 ± 8.1 pA/pF, n = 41; *p* < 0.01; A246^IC^: 50.7 ± 5.4, n = 10; *p* < 0.001) (**Figure 3C**). Hence, the ratio between Nav and Kv currents (INav/IKv) was significantly larger for both isogenic controls compared to their respective fAD lines (*p* < 0.05) (**Figure 3D**), highlighting a potential imbalance between inward and outward currents as the underlying cause of reduced excitability in *PSEN1* fAD iNs.

**Figure 3.**
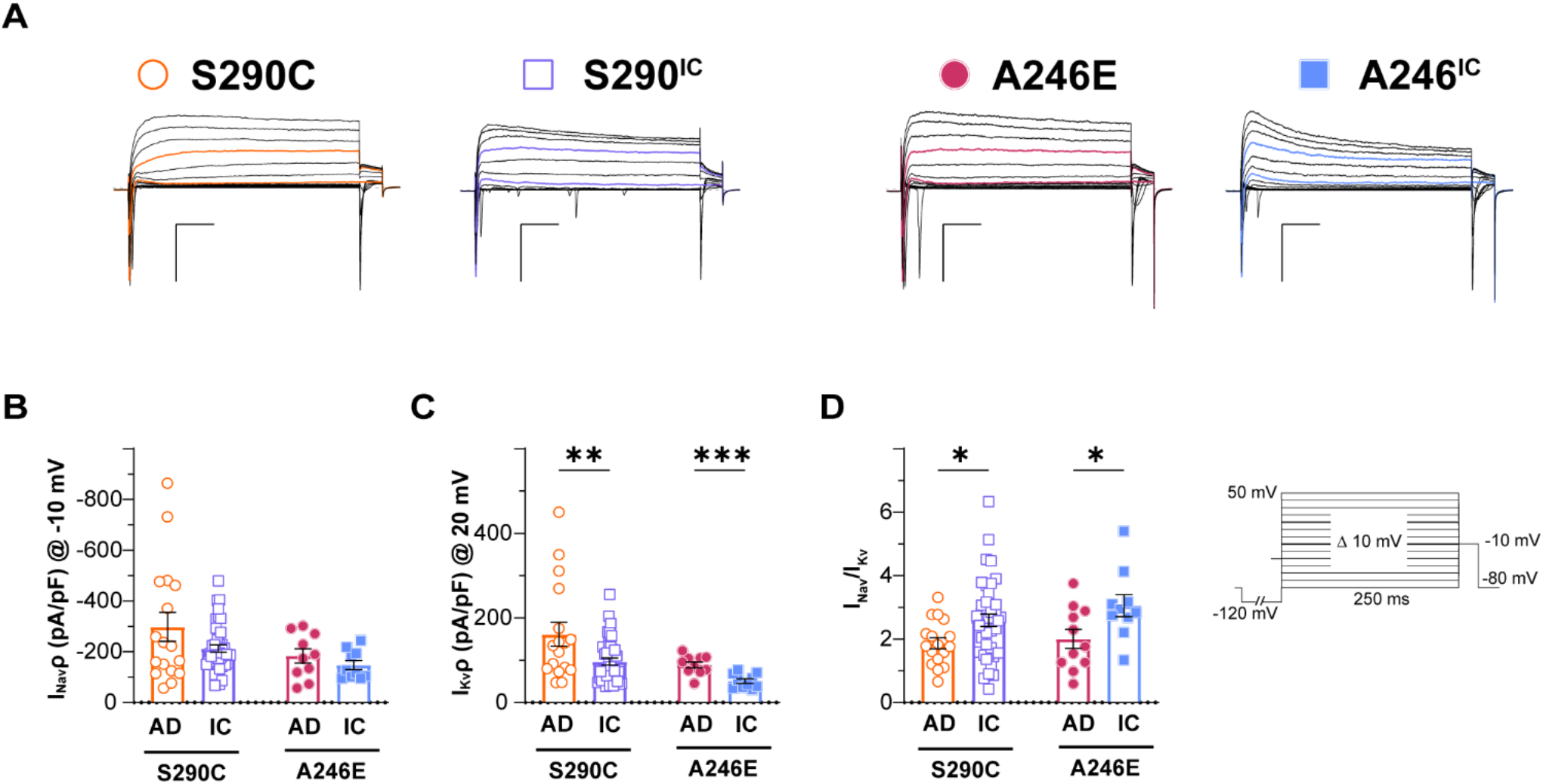
Active membrane properties of neurons derived from AD patients with a *PSEN1* mutation and their isogenic controls. Whole cell patch-clamp recordings were performed on day 21-35 NGN2 iNs derived from S290C (PSEN1 S290C) and A246E (PSEN1 A246E) and their respective CRISPR corrected controls, S290IC and A246IC cell lines. (**A**) Representative traces of Nav and Kv currents elicited by 250 ms square pulses from −80 mV to 50 mV in 10 mV steps (Vh −80 mV, 0.1 Hz). A 250 ms pre-pulse to −120 mV was used to minimize cumulative inactivation (inset). Nav (**B**) and Kv (**B**) current density quantification. (**C**) Nav to Kv current ratio. Data is presented as the mean ± SEM. Each data point represents an individual cell (n = 10 – 41), from 3 independent differentiations. Data was analysed using multiple t-tests with Holm-Sidak for multiple comparisons where \**p* < 0.05, \*\**p* < 0.01, \*\*\**p* < 0.001 and \*\*\*\**p* < 0.0001. **Abbreviations**: AD – Alzheimer’s disease, IC – isogenic control.

## 4 Discussion

This study investigated early excitability changes in fAD using iPSC-derived neurons from patients bearing *PSEN1* mutations. These neurons showed reduced firing responses suggestive of altered ionic homeostasis. Action potential properties in neurons bearing S290C or A246E mutations diverged, indicating varied disrupted pathways. Understanding these early cellular and molecular changes may shed light on cause and progression in fAD.

There have been over 300 mutations in the *PSEN1* gene identified with varying pathogenicity and effects on neuropathology and molecular pathways. This study has provided evidence of common intrinsic excitability changes that are altered in neurons with a *PSEN1* S290C or A246E mutations. The S290C mutation occurs due to a missense mutation in the splice acceptor site on the boundary of intron 8 and exon 9, resulting in the skipping of exon 9, which leads to impaired APP cleavage and Aβ processing (Rovelet-Lecrux *et al*. 2015; Steiner *et al*. 1999). The deletion of exon 9 impairs cellular functions including calcium influx in hippocampal neurons (Skobeleva *et al*. 2022), altered mitochondrial metabolism, calcium homeostasis and inflammation in astrocytes (Oksanen *et al*. 2017), impaired endocytosis in neurons (Woodruff *et al*. 2016) and lipid metabolism (Landman *et al*. 2006). In this research, S290C iNs exhibited reduced capacitance, indicative of smaller cell sizes or reduced neurite extensions and a decreased Nav/Kv ratio, likely underscoring the smaller AP area/rise parameters and overall impaired neuronal excitability. The A246E mutation is also pathogenic and occurs in exon 7 of *PSEN1* (Sherrington *et al*. 1995), impeding proper APP cleavage and resulting in toxic amyloid peptide generation (Mahairaki *et al*. 2014; Yang *et al*. 2017). In a cellular context the A246E mutation results in impaired microglial differentiation (Aubert *et al*. 2023), premature neuronal differentiation (Yang *et al*. 2017), altered astrocyte metabolic function and inflammatory activation (Elsworthy *et al*. 2023) and increased neuronal susceptibility to Aβ (Armijo *et al*. 2017). In this study, iNs with the *PSEN1* A246E mutation showed decreased hyperpolarizing responses which together with an Nav/Kv imbalance would require stronger current injection (rheobase) for AP firing.

The observed reduced neuronal excitability in iNs in this study is consistent with previous findings on *PSEN1* fAD mutations. For instance, cortical organoids with an L345F mutation displayed reduced extracellular activity measured by multi-electrode array analysis compared to its isogenic control, likely due to altered notch signaling (Hurley *et al*. 2023). The A246E cell line used in this study, was demonstrated to have a deficiency in Notch1 in iNs, resulting in susceptibility to ferroptosis (Greenough *et al*. 2022). In APP_SWE_/PSEN1(dE9) transgenic mice, hippocampal neurons exhibited increased rheobase, decreased action potential frequency at lower current injections (30 and 50 pA), and increased frequency at 100 pA (Chen *et al*. 2021). Time-dependent changes in spike frequency of CA1 pyramidal neurons were observed in the APP_SWE_/PSEN1(dE9) transgenic mouse model, with reduced spike counts compared to wild-type at 1 month of age, no differences at 4 months and increased spikes at 10 months, suggesting excitability is altered with disease progression (Vitale *et al*. 2021). Future studies should investigate the impact of the S290C mutation on Notch signaling as many mutations have been linked to impaired cleavage of Notch1, resulting in impaired neurogenesis or premature differentiation, which may be an explanation for the altered excitability changes. Zhang *et al*. (2020) showed the *PSEN1* S169 deletion mutation, induced AD pathology and cognitive deficits in a notch signaling independent pathway, suggesting different *PSEN1* mutations may contribute to neuronal deficits via distinct mechanisms. This phenomenon may also explain specific impacts on action potential properties between lines. Common to both mutations was enhanced Kv conductance, likely implicated in hypoexcitability. Kv1 channels are known to regulate the repression of intrinsic excitability and synaptic transmission (Thouta *et al*. 2021; Colasante *et al*. 2020) and are upregulated in AD proteomic datasets (Askenazi *et al*. 2023). Nevertheless, the vast collection of potassium channels expressed in neurons and glia warrant future work identifying Kv channel expression changes in fAD *PSEN1* mutant cell lines as well as investigating the effects of introducing the same *PSEN1* mutations into otherwise healthy cell lines to determine direct causal links between presenilin-1 dysfunction and Kv channel dysregulation.

In a reductionist model, this study revealed a common hypoexcitability phenotype in iNs generated from fAD patients with *PSEN1* mutations in the absence of Aβ and tau disease pathology (Targa Dias Anastacio *et al*. 2024) and supporting cell types. Future research may employ genome editing to create cell lines with *PSEN1* mutations of varying pathogenicity to establish a potential link between neuronal excitability and early onset-AD progression. In animal models, Aß appears to induce neuronal hyperexcitability), while soluble mutant tau suppresses neural activity in the rTg4510 and P301S tau transgenic mouse models (Busche *et al*. 2019; Marinković *et al*. 2019; Menkes-Caspi *et al*. 2015). Ghatak *et al*. (2019) reported hyperexcitability in co-cultures of astrocytes and iPSC-derived neurons from fAD patients. It is hypothesised that early stages of AD results in Aß-dependent hyperexcitability, preceding the onset of tau-dependent hypoactive neural circuits (Harris *et al*. 2020). Differences in neuronal maturity, Aß and tau generation between cell lines and co-culture may influence neural circuits *in vitro*. To determine the interplay between *PSEN1* mutations, AD pathology and neuronal excitability, simplistic 2D models may not be sufficient and would require more advanced systems such as 3D cerebral organoids matured long-term. It is also important to consider how support cells such as oligodendrocytes, astrocytes and microglia, influence the disruption to neuronal activity and transition between hypo- and hyperexcitability.

In summary, this study demonstrates that neurons derived from AD patients with *PSEN1* mutations exhibit reduced firing activity and altered electrophysiological properties. Mechanistic understanding of the early changes disrupted in AD will provide insight into the driving forces of neurodegeneration and proved novel avenues for intervention to slow this devastating disease.

## 7 Author Contributions

SM: conceptualisation, data curation, formal analysis, investigation, methodology, project administration, visualisation, writing (original draft), writing (review and editing). R.K.F-U: conceptualisation, investigation, methodology, resources, validation, visualisation, writing (review and editing). A.J.H: data curation, investigation, methodology, writing (review and editing). M.C.C-S, H.T.D.A., R.B., T.B., C.T., S.S.M., M.E., P.K., L.L., K.S., P.S.: cell line generation and characterisation. M.D: conceptualisation, funding acquisition, resources, supervision, writing (review and editing). L.O: conceptualisation, funding acquisition, resources, supervision, writing (review and editing).

## 9 Acknowledgments

The authors would like to thank the donors for their contribution to make this research possible. The authors would like to thank D.J. Adams for comments on the manuscript, providing the facilities, resources, and support for R.K.F-U (ARC Grant DP210102405 awarded to D.J. Adams).

## 10. Data Availability Statement

The datasets generated in this study can be requested from the corresponding authors upon reasonable request.

