## Supplementary material for "Alzheimer’s disease induced neurons bearing *PSEN1* mutations exhibit reduced excitability"

**Supplementary methods**

1. **Cell line generation and characterisation**
   1.1. **Reprogramming**

Dermal fibroblasts were collected from a 43 year old male diagnosed with Alzheimer’s disease harbouring a *PSEN1*^S290C^ mutation and had an The donor had an *APOE* ɛ3/ɛ3 genotype. Fibroblasts were cultured in fibroblast medium (DMEM/F12 supplemented with 10% foetal bovine serum (Interpath, #SFBS-F), 1 x L-Gluatmine (Life Technolgies, #25030) and 1x PenStep (Sigma Aldrich, P4333) at 37°C, 5% CO_2_ as described in (Bax *et al.* 2019). Induced pluripotent stem cells (iPSCs) were generated through reprogramming of fibroblasts using the microRNA-enhanced mRNA reprogramming kit (Stemgent, #00-0073, #00-0071) as per manufacturer’s guidelines. Colonies were manually selected and iPSCs were cultured in mTESR1 (Stem Cell Technologies, #85850), with daily media changes and passaged weekly. The cell line is referred to as S290C for simplicity (cell line ID: UOWi009-A <https://hpscreg.eu/cell-line/UOWi009-A>).

1.2. **CRISPR-gene correction**

The S290C cell line was CRISPR-gene corrected through the generation of a clonally derived cell line was generated by delivering components of CRISPR/SpCas9 system via plasmid - pSpCas9(BB)-2A-Puro (PX459) V2.0 (Addgene Plasmid #62988) together with 67nt long single-stranded (ss)DNA template (IDT) (10uM). Both plasmid and ssDNA were delivered by electroporation using Amaxa 4D Nucleofector using modified Lonza iPSC nucleofection protocol for 100ul reaction using solution P3 and program CA137. Transfected cells were selected on Puromycin (0.5ug/ml) for the next 24 hours. Edited cells were fed for the next 4 days with mTeSR™ medium (STEMCELL Technologies) containing CloneR™ supplement (STEMCELL Technologies), after which they were expanded only in mTeSR™ medium (STEMCELL Technologies). Clonally derived cell line was generated from edited mixed pool by limiting dilution in 96 well plate. Genomic target for SpCas9 nuclease is sequence 5’ GTCTTTCCCAACATCAACAA 3’ while PAM sequence is 3 nt sequence TGG. PCR primers flanking targeted genomic region are - Forward - 5’-AGTGGAATCTTGCTGGCCTG-3’ and Reverse 5’-GAGCTTCCGGGTCTCCTTCT-3’ producing amplicon length of 362bp. The CRISPR corrected isogenic control is referred to as S290^IC^ (UOWi009-A-1 <https://hpscreg.eu/cell-line/UOWi009-A-1>).

1.3. **Karyotyping**

Karyotyping of the S290C line was performed by StemCore at the University of Queensland, and the isogenic control was karyotyped at Monash Health. Cells were assessed at a resolution of 400 bands per haploid set.

1.4. **Sequencing**

Sequencing was performed using primers targeting *PSEN1* (**Table S1**) as previously described in Muñoz *et al.* (2018) and Balez *et al.* (2020).

1.5. **RT-qPCR**

Total RNA was extracted from cultures using TRIzol reagent (Life Technolgies, #15596-026), genomic DNA was removed using Turbo DNAse kit (Thermo Fisher Scientific, # AM1907) and cDNA was generated using the Tetro cDNA synthesis kit (Bioline, #BIO-65043). Gene expression was analysed using SensiFast SYBR green mastermix (Bioline, #BIO-98005) on the QuantStudio 5 (Applied Biosystems). Relative gene expression was calculated by normalizing to the housekeeper genes, *GAPDH* and *HPRT1*.

1.6. **Immunocytochemistry**

iPSCs were fixed with 4% paraformaldehyde for 15 min at RT and permeabilised with 0.3% Triton X-100 (Sigma-Aldrich, #T9284) for 15 min at RT. Cells were blocked with 10% goat serum in PBS for 1 h at RT. Cells were incubated with primary antibodies overnight at 4°C (Table 2), followed by secondary antibody incubation for 1 h at RT (**Table S1**). Nuclei were stained with Reddot 2 (1:200, Biotum, #40061-1) for UOWi009-A and Hoechst 3342 (1 µg/mL) for UOWi009-A-1 for 10 min at RT. Images were acquired on the Leica SP5 or SP8 confocal microscope.

1.7. **Scorecard**

iPSCs were differentiated into mesoderm and endoderm lineages using the STEMdiff mesoderm induction medium (Stem Cell Technologies, # 05220) and STEMdiff definitive endoderm kit (Stem Cell Technologies, # 05110), respectively. Induction of ectodermal differentiation was completed as described in Muñoz *et al.* (2018) and Balez *et al.* (2020). From each lineage differentiation, cDNA was pooled at a 1:1:1 ratio with 1 µg analyzed with the Taqman hPSC scorecard (Thermo Fisher Scientific, # A15871).

1.8. **STR analysis**

STR profiling of 18 locations was performed at the Garvan Molecule Genetics Institute (Darlinghurst, Australia).

**Table S1. Reagent details.**

| **Antibodies used for immunocytochemistry** | | | |
| --- | --- | --- | --- |
|  | **Antibody** | **Dilution** | **Cat # and RRID** |
| Pluripotency Markers | Anti-Human SSEA-4 | 1:200 | Abcam, ab16287, RRID: AB_778073 |
|  | Anti-human OCT4 | 1:500 | Stem Cell Technologies, 01550, RRID: AB_1118539 |
| Secondary Antibodies | Alexa Fluor 488 Goat anti-mouse IgG (H + L) | 1:1000 | Thermo Fisher Scientific, A11001, RRID: AB_2534069 |
| **Site-specific nuclease** | | | |
| Nuclease information | *Streptococcus pyogenes (Sp)* | Sequence from Addgene Plasmid #62988 | |
| Delivery method | Transfection | Addgene Plasmid (#62988) together with 67nt long single-stranded (ss)DNA template (IDT) | |
| Selection/enrichment strategy | Puromycin (0.5ug/ml) | Post transfection selection with puromycin (0.5ug/ml) for 24 hours | |
| **Primers and oligonucleotides** | | | |
|  | **Target** | **Forward/Reverse primer (5′-3′)** | |
| Sequencing | *PSEN1* | Forward: GTGTGGAGAAATGATGGC  Reverse: CTTATCTGTGTATTTACTGGGC | |
| Pluripotency markers (qPCR) | *NANOG* | Forward: CCAGAACCAGAGAATGAAATC  Reverse: TGGTGGTAGGAAGAGTAAAG | |
| Pluripotency markers (qPCR) | *POU5F1* | Forward: GATCACCCTGGGATATACAC  Reverse: GCTTTGCATATCTCCTGAAG | |
| House-Keeping Genes (qPCR) | *GAPDH* | Forward: GAGCACAAGAGGAAGAGAGAGACCC  Reverse: GTTGAGCACAGGGTACTTTATTGATGGTACATG | |
|  | *HPRT1* | Forward: TGACACTGGCAAAACAATGCA  Reverse: GGTCCTTTTCACCAGCAAGCT | |
| gRNA oligonucleotide sequence | 5’GTCTTTCCCAACATCAACAA 3’ | | |

**Supplementary results**

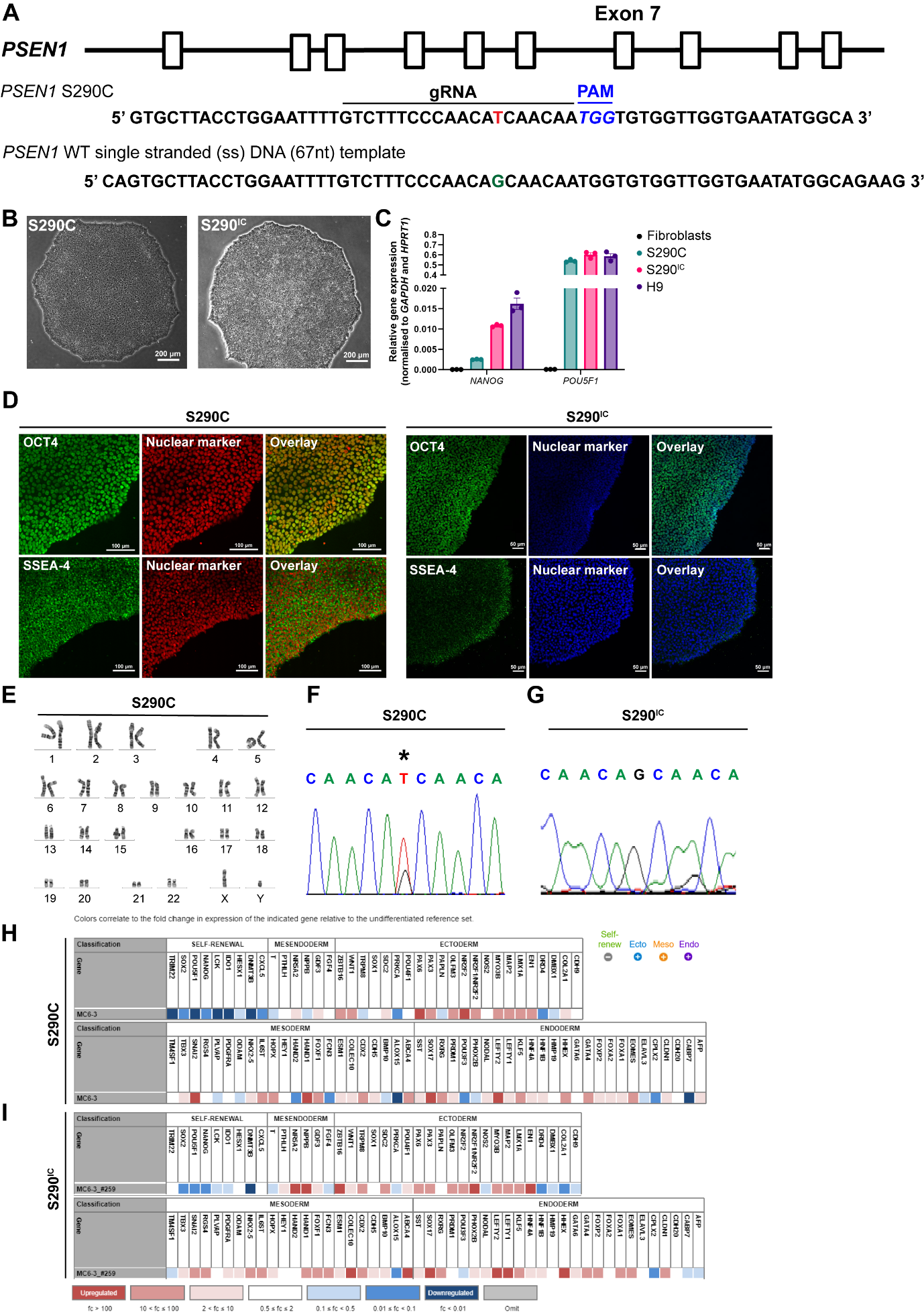

**Figure S1. Generation of iPSCs from a patient with a *PSEN1* S290C mutation and an isogenic control.** Dermal fibroblasts were collected from a 43 year old male bearing a *PSEN1*S290C mutation diagnosed with Alzheimer’s disease at the time of collection. **(A)** CRISPR was performed with pSpCas9(BB)-2A-Puro (PX459) V2.0 (Addgene Plasmid #62988) and single -stranded (ss) DNA template where SpCas9 and gRNA targeting *PSEN1*. **(B)** Representative images of colonies from both cell lines exhibiting normal morphology. **(C)** Expression of pluripotency marker genes POU5F1 (encodes OCT4) and NANOG were upregulated compared to fibroblasts, and were similar levels to the H9, human embryonic stem cell line as analysed by RT-qPCR. **(D)** Pluripotent markers OCT4 and SSEA-4 were confirmed at the protein level though immunocytochemistry. **(E)** Karyotyping revealed no abnormalities present in both the S290C. Sequencing identified that the mutation was present in the S290C line **(F)** and successfully corrected in S290^IC^ **(G)**. The capacity of iPSC lines to differentiate into all three germ layers was validated by a Taqman scorecard for S290C **(H)** and S290^IC^ **(I)**.

**Table S2. Whole-cell patch clamp data.**

|  | **S290C** | | | **S290^IC^** | | | **A246E** | | | **A246^IC^** | | |
| --- | --- | --- | --- | --- | --- | --- | --- | --- | --- | --- | --- | --- |
|  | *Mean* | *SEM* | *n* | *Mean* | *SEM* | *n* | *Mean* | *SEM* | *n* | *Mean* | *SEM* | *n* |
| **RMP** | -48.4 | 4.6 | 13 | -49.8 | 1.4 | 21 | -63.3 | 2.7 | 8 | -46.6 | 2.1 | 9 |
| **Capacitance** | 13.0 | 0.8 | 17 | 15.5 | 0.6 | 41 | 14.2 | 1.4 | 11 | 13.8 | 0.9 | 10 |
| **Rheobase** | 41.5 | 9.8 | 13 | 19.5 | 3.0 | 21 | 66.3 | 9.8 | 8 | 22.2 | 4.6 | 9 |
| **#APs @2xrheo** | 2.0 | 0.7 | 12 | 6.6 | 0.8 | 21 | 1.4 | 0.4 | 8 | 4.0 | 1.0 | 9 |
| **-150 pV hype** | -82.8 | 11.4 | 12 | -101.4 | 5.8 | 21 | -85.8 | 12.8 | 8 | -141.0 | 9.5 | 9 |
| **Max decay slope** | -20.8 | 2.8 | 13 | -17.0 | 1.1 | 21 | -28.7 | 3.8 | 8 | -11.9 | 1.4 | 9 |
| **AP half-width** | 7.5 | 2.0 | 13 | 5.3 | 0.3 | 21 | 4.2 | 0.5 | 8 | 8.0 | 1.2 | 9 |
| **AP area** | 2229.0 | 423.5 | 13 | 12036.2 | 1390.3 | 21 | 8518.6 | 3015.7 | 8 | 11285.8 | 2004.2 | 11 |
| **Peak amplitude** | 74.2 | 8.2 | 13 | 83.5 | 3.0 | 21 | 99.7 | 3.3 | 8 | 72.9 | 3.7 | 9 |
| **Rise slope** | 25.6 | 4.1 | 13 | 59.3 | 9.4 | 21 | 42.6 | 6.2 | 7 | 26.6 | 4.7 | 9 |
| **Rise time** | 1.3 | 0.2 | 13 | 0.6 | 0.1 | 20 | 0.8 | 0.2 | 7 | 1.2 | 0.2 | 9 |
| **Kv** | 161.5 | 28.1 | 17 | 97.0 | 8.1 | 41 | 89.2 | 7.1 | 10 | 50.7 | 5.4 | 10 |
| **Nav** | -298.4 | 56.9 | 17 | -213.1 | 14.7 | 41 | -184.3 | 28.0 | 10 | -148.0 | 17.8 | 10 |
| **Ina/IK** | 1.9 | 0.2 | 17 | 2.6 | 0.2 | 41 | 2.0 | 0.3 | 11 | 3.1 | 0.3 | 10 |

Balez, R., Berg, T., Bax, M. et al. (2020) The mRNA-based reprogramming of fibroblasts from a SOD1E101G familial amyotrophic lateral sclerosis patient to induced pluripotent stem cell line UOWi007. *Stem Cell Research* **42,** 101701.

Bax, M., Balez, R., Muñoz, S. S. et al. (2019) Generation and characterization of a human induced pluripotent stem cell line UOWi005-A from dermal fibroblasts derived from a CCNFS621G familial amyotrophic lateral sclerosis patient using mRNA reprogramming. *Stem Cell Research* **40,** 101530.

Muñoz, S. S., Balez, R., Berg, T. et al. (2018) Generation and characterization of human induced pluripotent stem cell lines from a familial Alzheimer's disease PSEN1 A246E patient and a non-demented family member bearing wild-type PSEN1. *Stem cell research* **31,** 227-230.
